# Parasite-probiotic interactions in the gut: *Bacillus* sp. and *Enterococcus faecium* regulate type-2 inflammatory responses and modify the gut microbiota of pigs during helminth infection

**DOI:** 10.1101/2021.09.01.458597

**Authors:** Laura J. Myhill, Sophie Stolzenbach, Helena Mejer, Lukasz Krych, Simon R. Jakobsen, Witold Kot, Kerstin Skovgaard, Nuria Canibe, Peter Nejsum, Dennis S. Nielsen, Stig M. Thamsborg, Andrew R. Williams

## Abstract

Dietary probiotics may enhance gut health by directly competing with pathogenic agents and through immunostimulatory effects. These properties are recognized in the context of bacterial and viral pathogens, but less is known about interactions with eukaryotic pathogens such as parasitic worms (helminths). In this study we investigated whether two probiotic mixtures (comprised of *Bacillus amyloliquefaciens, B. subtilis*, and *Enterococcus faecium* [BBE], or *Lactobacillus rhamnosus* LGG and *Bifidobacterium animalis* subspecies *Lactis* Bb12 [LB]) could modulate helminth infection kinetics as well as the gut microbiome and intestinal immune responses in pigs infected with the nodular worm *Oesophagostomum dentatum*. We observed that neither probiotic mixture influenced helminth infection levels. BBE, and to a lesser extent LB, changed the alpha- and beta-diversity indices of the colon and faecal microbiota, notably including an enrichment of faecal *Bifidobacterium* spp. by BBE. However, these effects were muted by concurrent *O. dentatum* infection. BBE (but not LB) significantly attenuated the *O. dentatum-induced* upregulation of genes involved in type-2 inflammation and restored normal lymphocyte ratios in the ileo-caecal lymph nodes that were altered by infection. Moreover, inflammatory cytokine release from blood mononuclear cells and intestinal lymphocytes was diminished by BBE. Collectively, our data suggest that selected probiotic mixtures can play a role in maintaining immune homeostasis during type 2-biased inflammation. In addition, potentially beneficial changes in the microbiome induced by dietary probiotics may be counteracted by helminths, highlighting the complex inter-relationships that potentially exist between probiotic bacteria and intestinal parasites.

## Introduction

The mammalian gut environment is maintained in a complex homeostasis encompassing interactions between dietary compounds, the commensal gut microbiota (GM) and the mucosal immune system. Dysregulation of this balanced ecosystem can lead to increased susceptibility to pathogen infection and chronic inflammation, and is a major source of disease and morbidity in humans and decreased productivity in livestock. To this end, dietary supplementation with probiotic bacteria has gained increasing attention as a safe method to maintain intestinal homeostasis, subsequently improving gut health. Beneficial effects of probiotics are strain-specific and dose-dependent, and can be achieved by modulating intestinal motility and barrier function, outcompeting enteropathogens, or by modifying the composition of host GM, subsequently affecting host mucosal immune responses ^1, 2^.

Pigs are a key species in the food production industry and also serve as an important model for human biomedical research due to similarities in gastrointestinal physiology and microbiota composition ^3^. Supplementation of pig diets with probiotics has revealed beneficial effects such as improved growth, carcass quality, and enhanced host protective responses against different pathogens, with pronounced efficiency at reducing bacterial load of enterotoxigenic *Escherichia coli* (F4) in weaned piglets ^4–8^. Additional studies against eukaryotic pathogens have also reported beneficial effects of probiotics. For example, *in vitro* and murine models of *Giardia* infection have shown that *Lactobacillus* spp. and *Enterococcus faecium* can eliminate infection and reinforce host immune responses ^9–11^.

Parasitic worms (helminths) are among the most widespread gut pathogens, infecting more than a billion humans worldwide and being commonly found in nearly all farmed livestock ^12, 13^. Infection can result in marked immunopathology and a reduction in mucosal barrier function and poses a significant risk to health and productivity ^14, 15^. Moreover, mucosal-dwelling helminths induce strongly polarized T helper (Th) type-2 immunity and thus serve as a useful model for Th2-mediated intestinal immune responses, such as those elicited by food allergens ^16^. Studies on the trilateral interactions between parasites, the GM and the immune system may therefore shed light on the role of gut bacteria in regulating host-parasite and immune interactions at mucosal barrier surfaces. Several studies have reported that feeding prebiotic dietary fibres (e.g. inulin) or administration of microbial metabolites (short-chain fatty acids (SCFA) or lactic acid) can strongly influence infection dynamics and immune responses induced by the large intestinal-dwelling parasites *Trichuris suis* (porcine whipworm) and *Oesophagostomum dentatum* (porcine nodular worm) ^17–19^. These effects are thought to be mediated by GM changes in the caecum and colon ^19, 20^, as inulin is known to increase the abundance of microbes such as *Lactobacillaceae* and *Bifidobacterium* during helminth infection ^19^.

Reports on the effects of dietary probiotic supplementation during helminth infection are limited, and whether probiotics can modulate helminth infection and associated inflammatory and immunopathological changes in the large intestine, as appears to be the case with inulin and other prebiotics, remains unknown. Supplementation with *Bifidobacterium animalis* subspecies *Lactis* Bb12 was shown to modulate mucosal immune responses and enhance jejunal barrier function in pigs infected with *Ascaris suum*, whilst *Lactobacillus rhamnosus* LGG intake supressed the development of type-2 related immune responses in the tracheal-bronchial lymph nodes of *A. suum*-infected pigs ^22^. Thus, probiotic bacteria may exert immunomodulatory effects in the context of type-2 immune function. In light of this, porcine models of helminth infection may represent a valuable model for studying the interactions between probiotic bacteria and gut pathogens, and assessing if probiotics have potential as health-promoting dietary additives that can prevent or alleviate the effects of enteric helminth infection.

Here, we investigated the effects of two different probiotic mixtures on *O. dentatum* establishment and infection dynamics in pigs. In addition, we explored the interactions between these probiotics and infection on GM composition throughout the intestinal tract, as well as peripheral and local mucosal immune responses. We show that a dynamic relationship exists between probiotic supplementation, the GM and the immune system during helminth infection, which may have significant implications for our understanding of the regulation of type-2 inflammation in mucosal tissues, and for the application of probiotics for prevention or control of intestinal diseases.

## Results

### Effects of probiotics on the intestinal environment and *O. dentatum* infection levels

Pigs (n=48) were divided into three groups (**Supplementary Figure 1**). 16 pigs received only the basal control diet (based on ground barley and soybean meal) throughout the study, 16 pigs received the basal diet supplemented with a mixture of *Bacillus amyloliquefaciens B. subtilis*, and *Enterococcus faecium* (hereafter BBE), and 16 pigs received the basal diet supplemented with a mixture of *Lactobacillus rhamnosus* LGG and *Bifidobacterium animalis* subsp*. Lactis* BB-12 (hereafter LB). The BBE mixture was selected based on its development specifically to improve gut health in pigs, whilst LB was chosen as it contained two well-studied probiotic strains that have previous been shown to induce immunomodulatory activity in pigs ^21, 23^. Within each dietary group, following a 14 day acclimatization period, half the pigs (n=8) were either trickle-infected throughout the study with *O. dentatum* larvae (n=24), or remained uninfected (n=24).

To explore the effects of probiotics on the response to helminth infection, we quantified the effect of probiotic supplementation on intestinal physicochemical parameters and parasite establishment and development. We first assessed the concentrations of SCFA and D-lactic acid in the proximal colon (**Figure 1A**), with a separate analysis conducted for the two different probiotic-supplemented groups, relative to those with no supplementation. Acetic and propionic acid concentrations were unaffected by either infection or probiotic supplementation. *O. dentatum* infection significantly increased n-Butyric acid levels (*p* < 0.05) in pigs fed either the control diet alone or in those supplemented with BBE. However, there was no effect of *O. dentatum* when analysing LB-supplemented pigs, indicating that the effect of infection varied according to specific probiotic intake (**Figure 1A**). Total SCFA levels were not different between any of the groups. In contrast, D-lactic acid levels were significantly increased by LB supplementation, and tended also to be increased by BBE supplementation (*p* = 0.08), independently of infection (**Figure 1A**). Neither probiotic supplementation nor infection influenced the pH in the jejunum or ileum (data not shown), or the caecum or proximal colon (Figure 1B). However, infection resulted in a lower pH in the distal colon (*p* < 0.05; **Figure 1B**).

**Figure 1.**
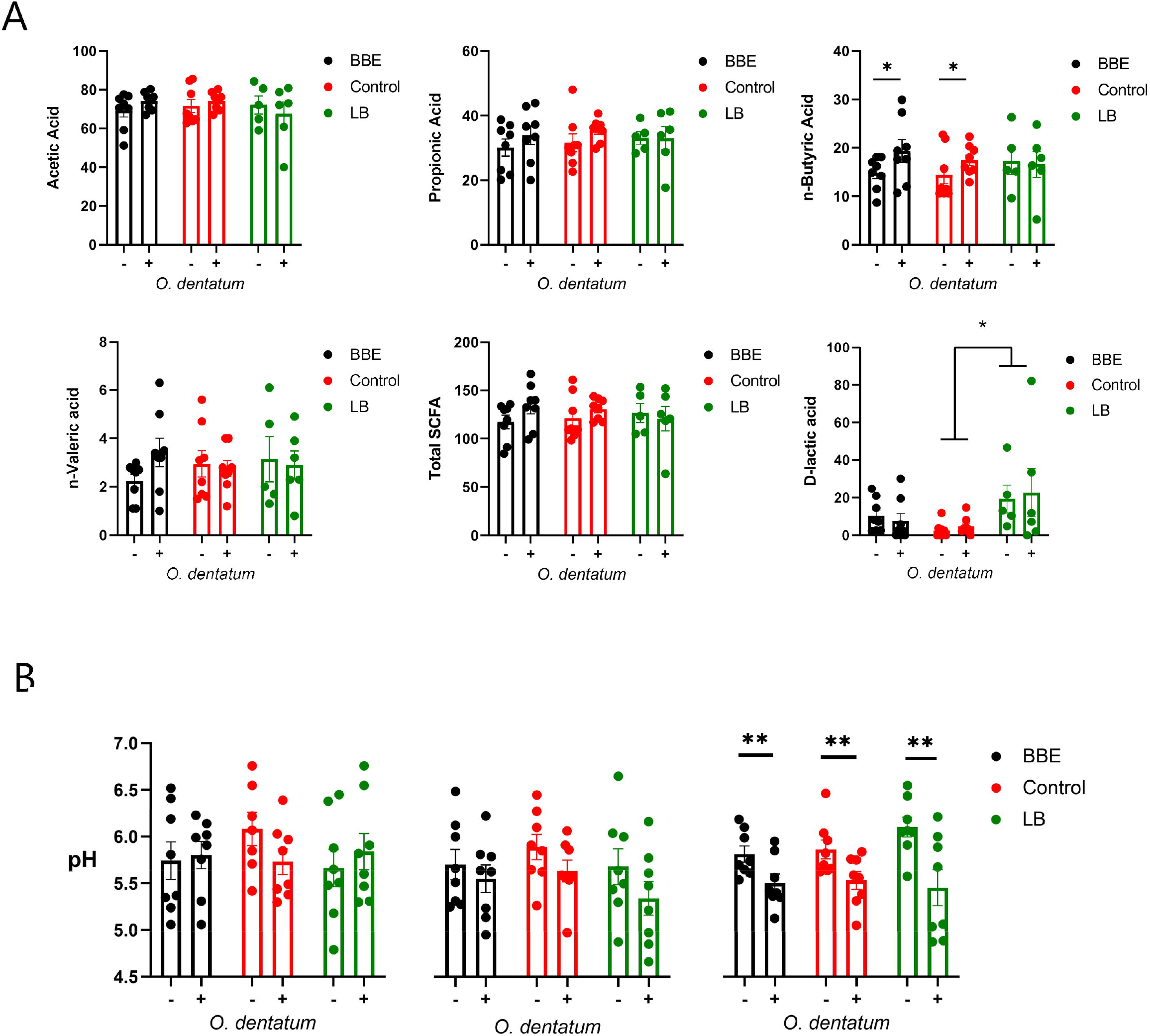
Effect of probiotics and *Oesophagostomum dentatum* infection on the intestinal environment. **(A)** Microbial metabolite (short-chain fatty acids and D-lactic acid) concentrations from proximal colon digesta after 28 days of *O. dentatum* infection, in pigs fed a control diet or a diet supplemented with either a mixture of *Enterococcus faecium* and *Bacillus* sp. (BBE), or LGG and Bb12 (LB). Metabolite concentrations are expressed in mmol/kg wet sample. **(B)** pH of digesta sampled throughout the intestinal tract. Statistical analysis was conducted separately for each probiotic treatment, using a GLM analysis comparing the effect of probiotic supplementation and infection (and their interaction) to the control-diet groups (no probiotics). Data presented as means ± SEM (\**p* ≤ 0.05, \*\*\**p* ≤ 0.005, by GLM). n=8 pigs per treatment group.

Supplementation with either of the probiotic mixtures did not significantly influence infection levels or parasite infection kinetics, with average worm numbers (adult and larval *O. dentatum*) of 15,843 ± 2,128 and 17,425 ± 2,185 (mean ± SEM) for pigs fed BBE and LB probiotics respectively, compared to 18,455 ± 2,598 for the control-fed group (**Figure 2**). Moreover, probiotic supplementation had no effect on worm length (data not shown) nor egg production; with similar eggs per gram faeces (EPG) scores observed for all diet treatment groups. In addition, no significant differences in body weight gain were observed between the dietary treatment/infection groups, with all pigs gaining weight consistently over the course of the experimental period (data not shown).

**Figure 2.**
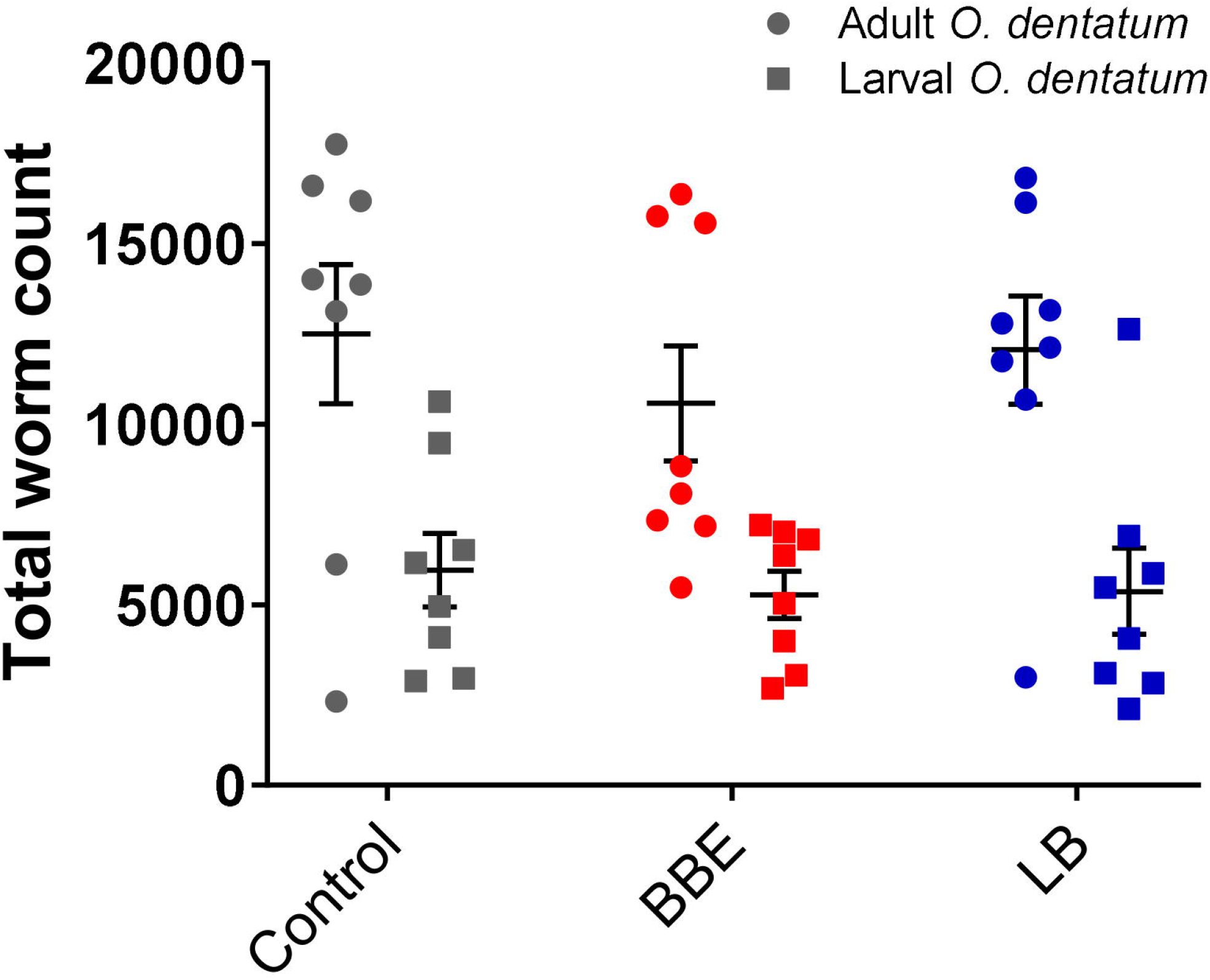
*Oesophagostomum dentatum* burden is not affected by probiotic treatment. *O. dentatum* adult and larval worm burdens, at day 28 post-infection in pigs fed a control diet or a diet supplemented with either a mixture of *Enterococcus faecium* and *Bacillus* sp. (BBE), or LGG and Bb12 (LB). Data presented as means ± SEM. n=8 pigs per treatment group.

### *O. dentatum* infection changes the response of the faecal microbiota to probiotic supplementation

To examine if the two probiotic mixtures and/or *O. dentatum* affected the composition of the prokaryotic GM, we conducted longitudinal sampling and analyses of faeces over the course of the study. Across the time period, α-diversity remained stable in pigs with no probiotic supplementation, regardless of whether they were infected with *O. dentatum* or not, with no significant differences in Faith phylogenetic diversity (PD) (**Figure 3A).** In contrast, in both uninfected and *O. dentatum*-infected pigs, BBE or LB supplementation tended to increase the Faith PD over time (indicative of a more diverse microbiota at the end of the study than at the start), (*p* = 0.065 for LB in infected pigs; *p* = 0.05 in other cases) (**Figure 3A**).

**Figure 3.**
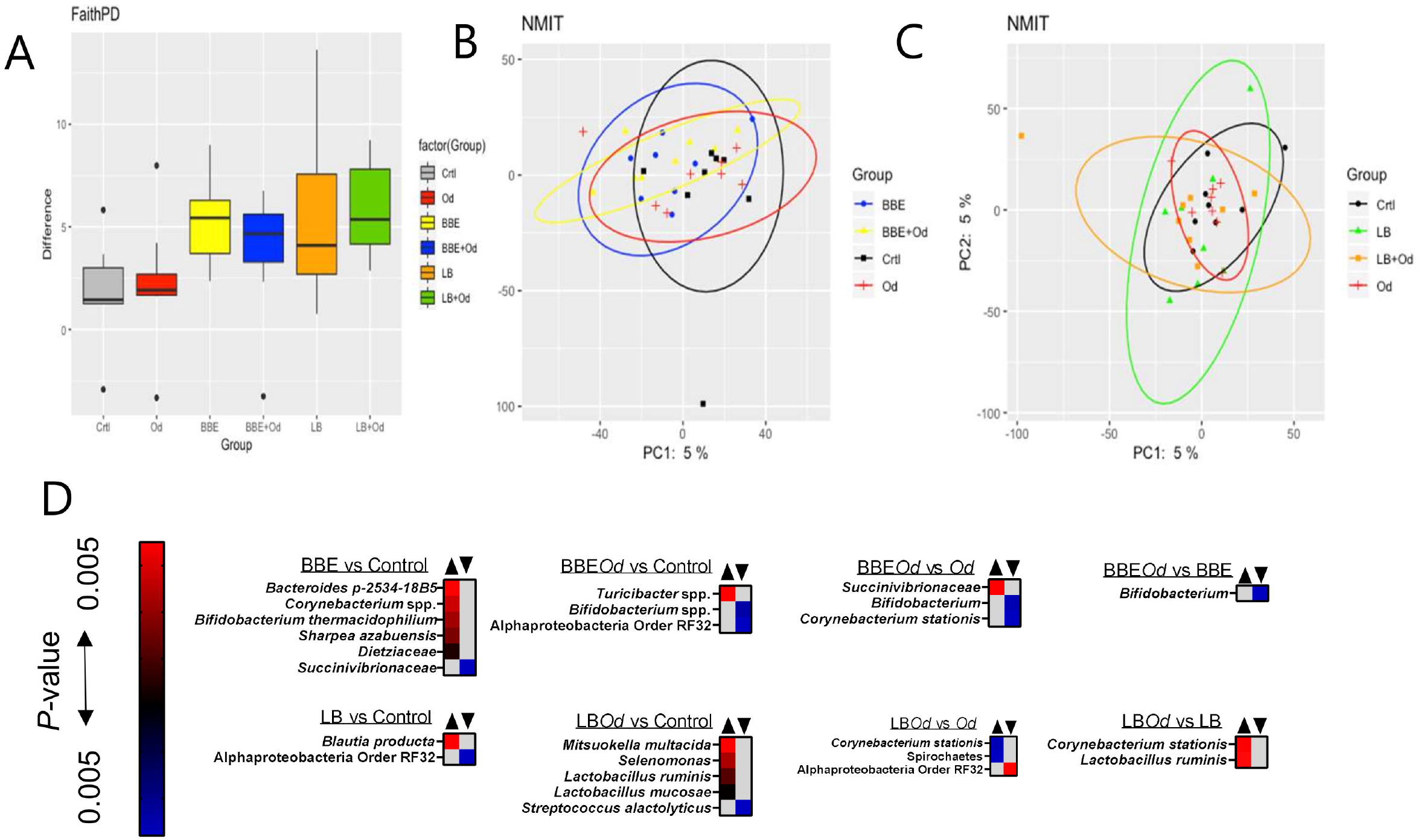
Probiotics modulate the faecal gut microbiota over time. **A)** Alpha-diversity (Faith PD) in faeces samples over time from −7 to day 28 post-infection (p.i.). Pigs were either uninfected or infected with *O. dentatum* (Od) and fed a control diet or a diet supplemented with either a mixture of *Enterococcus faecium* and *Bacillus* sp. (BBE), or LGG and Bb12 (LB). (**B-C**) NMIT PCoA showing effect of infection and diet in pigs fed BBE (**B**) or LB (**C**) from day −7 to day 28 p.i. **D)** Taxa where abundance was significantly altered in faeces across the course of the experiment as a result of infection or diet, as identified by Feature Volatility Analysis. n=8 pigs per treatment group.

There was also a significant shift in β-diversity in the faecal GM as a result of probiotic supplementation, but this was dependent on infection status. Non-parametric microbial interdependence testing (NMIT) indicated that infected pigs fed BBE differed in β-diversity from infected pigs without probiotic supplementation (*p* < 0.05; **Figure 3B**). However, this was not the case for uninfected pigs (*p* = 0.26; Figure 3B). A contrasting effect was observed for LB, where uninfected pigs fed LB diverged from uninfected pigs without probiotic supplementation (*p* < 0.05), yet infected pigs fed LB did not differ from infected pigs without LB (**Figure 3C**). In the absence of probiotic supplementation, infection did not influence β-diversity. Analyses on pooled data revealed a similar story, with both LB and BBE-fed pigs (independent of infection status) significantly diverging from control-fed pigs (*p* < 0.05), whereas infection status (independent of probiotic supplementation) had no effect (**Supplementary Figure 2**). Taken together, these data suggest that over the course of the seven week experiment, both BBE and LB probiotics induced modest but significant changes in the composition of the faecal microbiota, yet these probiotic-induced changes were further influenced by concurrent *O. dentatum* infection.

To explore which bacterial taxa were responsible for the divergence between probiotic-fed pigs and their respective controls without probiotics, Feature Volatility analysis was performed. Within uninfected pigs, six taxa were enriched in pigs receiving BBE compared to those that did not, most notably the *Bifidobacterium* genus, whilst a single family (*Succinivibrionaceae*) belonging to the Proteobacteria phylum decreased in abundance (**Figure 3D**). However, in infected pigs fed BBE, relative abundance of *Bifidobacterium* spp. was lower compared to infected pigs without BBE, indicating that the infection abrogated the probiotic-stimulated increase in *Bifidobacteria. Turicibacter* sp., a genus we have previously observed to be enriched in the colon of pigs infected with *Ascaris suum* ^24^, was elevated in infected pigs fed BBE compared to uninfected controls. Similarly, the effects of LB varied depending on infection status (**Figure 3D**). In uninfected pigs, only two taxa differed between LB-fed pigs and control-fed pigs without LB. In contrast, relative to the control group (uninfected pigs without probiotics), infected pigs fed LB had higher relative abundance of several members of the Firmicutes phylum including two *Lactobacillus* species, as well as *Mitsuokella multacida*, a putative butyrate producer and beneficial microbe ^25^. Collectively, these data suggest that BBE tended to enrich beneficial bacteria such as *Bifidobacterium* in faeces over the course of the experiment in uninfected pigs, but these effects were reversed in *O. dentatum*-infected pigs. Conversely, LB tended to enrich beneficial bacteria such as *Lactobacillus* more strongly in the faeces of *O. dentatum-infected* pigs than uninfected pigs. Thus, *O. dentatum* alone did not change the composition of the faecal microbiota over the course of the study, but instead modulated the effect of BBE and LB in two distinct ways, indicating a complex interaction between probiotics and the parasitic infection.

### Probiotics and *O. dentatum* infection interact to change the intestinal microbiota in a site-specific manner

We next investigated how infection and/or probiotics influenced the microbiota composition throughout the intestinal tract. Similarly to the longitudinal faecal samples, α-diversity (Faiths PD) was increased by both BBE and LB in comparison to control pigs, mainly in the distal colon, with a comparable effect in both infected and uninfected pigs (*p* = 0.093 for infected pigs fed LB; *p* < 0.05 for other comparisons; **Figure 4**). Notably, *O. dentatum* infection was also associated with increased α-diversity in the distal colon **(Figure 4; Supplementary Table 1**). Effects of infection and treatment were not as pronounced in the other gut segments (**Figure 4; Supplementary Table 1**).

**Figure 4.**
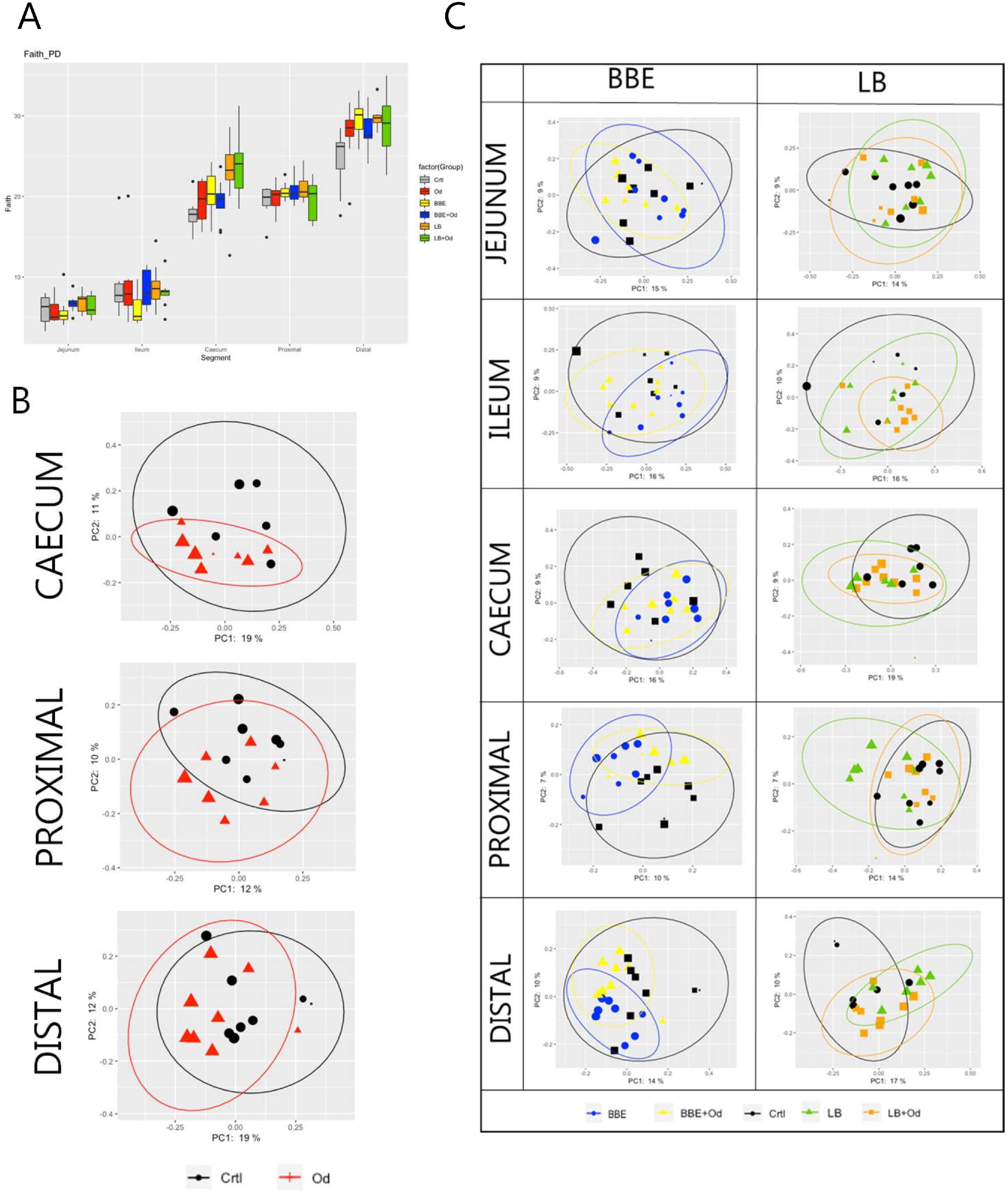
Probiotics and parasite infection modulate the gut microbiota in different gastrointestinal compartments. **A)** Alpha-diversity (Faith PD) in different gut segments at day 28 post-infection. Pigs were either uninfected or infected with *O. dentatum* (Od), and fed a control diet or a diet supplemented with a either mixture of *Enterococcus faecium* and *Bacillus* sp. (BBE), or LGG and Bb12 (LB). *P*-values are shown in Supplementary Table 1. n=8 pigs per treatment group. **B)** Unweighted PCoAs for pairwise comparisons of uninfected and *Oesophagostomum dentatum* (Od)-infected pigs fed only the control diet (no probiotics) in the caecum and proximal and distal colon. **C)** Unweighted PCoAs for pairwise comparisons of uninfected and *O. dentatum* (Od)-infected pigs fed only the control diet (no probiotics), or a diet supplemented with either a mixture of *Enterococcus faecium* and *Bacillus* sp. (BBE), or LGG and Bb12 (LB).

Analysis of unweighted Unifrac distance metrics showed that, in the absence of probiotic supplementation, the only intestinal site where *O. dentatum* infection significantly changed β-diversity, relative to uninfected pigs, was the proximal colon (the predilection site of the worms) (p < 0.05 by PERMANOVA; **Figure 4B**). β-diversity in the gut was also considerably altered by probiotic supplementation. Changes were primarily observed via unweighted Unifrac analysis, indicating that most differences were driven by low-abundance species. In uninfected pigs, BBE supplementation altered β-diversity compared to pigs without probiotic supplementation in the ileum, caecum and both proximal and distal colon (*p* = 0.096 for caecum, *p* < 0.05 for all other segments by PERMANOVA; **Figure 4C; Supplementary Table 2**). However, this effect was less evident when the BBE-supplemented pigs were infected with *O. dentatum*. In these animals, supplementation with BBE resulted in no significant difference in β-diversity in the ileum or caecum relative to control pigs (uninfected and without probiotics). Furthermore, lesser (albeit still significantly different) changes were observed in the colon between control pigs and infected pigs receiving BBE (**Figure 4C; Supplementary Table 2**). Thus, infection appeared to attenuate the BBE-induced changes in GM composition.

LB also tended to alter β-diversity in the jejunum and caecum, with similar changes in both uninfected and infected pigs (*p* < 0.1 by PERMANOVA; **Figure 4C; Supplementary Table 3**). LB had a stronger effect in the colon (both proximal and distal). Here, significant divergence was observed between control and LB-fed pigs, regardless of infection status (*p* < 0.05 by PERMANOVA). However, within LB-fed pigs, infected pigs were significantly diverged from uninfected pigs with infected pigs clustering closer to the control animals (*p* < 0.05 by PERMANOVA; **Figure 4C; Supplementary Table 3**), again indicating that infection tended to limit the modulatory effects of the probiotics on the GM.

We attempted to identify specific taxa responsible for the differences between treatment groups, however ANCOM analysis yielded no significant differences in any gut segment (*p* > 0.05). Thus, the changes in the GM community within the gut segments appeared to derive from the cumulative effect of subtle alterations across multiple taxa, rather than substantial alterations in the abundance of precise bacterial species.

### Both probiotics and *O. dentatum* infection influence peripheral and local immune function

We next assessed how probiotic supplementation modulated the development of the systemic and mucosal response to *O. dentatum* infection. Serum IgA and IgG_1_ antibody levels were measured weekly until day 28 p.i. All pigs were sero-negative for *O. dentatum* prior to study start at day 0. Infection with *O. dentatum* resulted in increased *O. dentatum-specific* antibody titres compared to uninfected pigs (**Figure 5A**). Both IgA and IgG1 antibody titre levels increased from day 7 through until day 28 p.i. There was a significant interaction between time and LB probiotics at day 21 p.i., whereby LB-fed infected pigs had higher IgA levels compared to the other infected groups (*p* < 0.005), however this difference was not apparent at other time points. BBE probiotics did not influence IgA titres, and there was no effect of probiotic supplementation on IgG1 titres.

**Figure 5.**
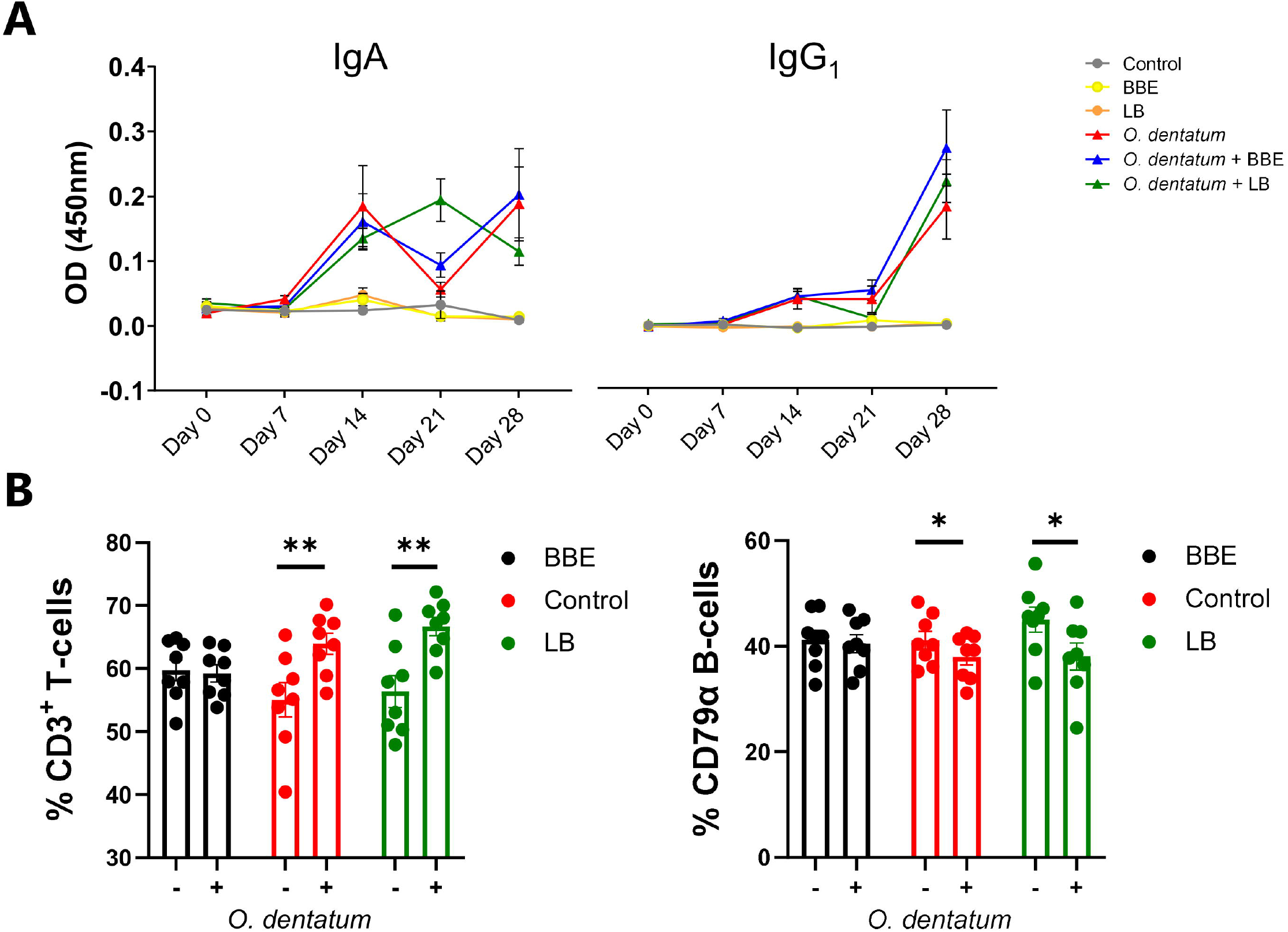
Systemic and peripheral immune responses elicited towards *Oesophagostomum dentatum* infection. **A)** *O. dentatum* specific IgA and IgG_1_ serum antibody production over the 28 days of infection in pigs fed a control diet or a diet supplemented with either a mixture of *Enterococcus faecium* and *Bacillus* sp. (BBE), or LGG and Bb12 (LB). **(B)** Flow cytometric analysis of ileo-caecal lymph node cells obtained at day 28 post-infection. % CD3+ T cells and % CD79α B-cells. Statistical analysis was conducted separately for each probiotic treatment, using a GLM analysis comparing the effect of probiotic supplementation and infection (and their interaction) to the control-diet groups (no probiotics). Data presented as means ± SEM (\**p* ≤ 0.05, \*\*\**p* ≤ 0.005, by GLM). n=8 pigs per treatment group.

Analysis of CLN lymphocyte populations revealed a significant interaction between BBE probiotic supplementation and *O. dentatum* infection. In control-fed pigs, infection increased the percentage of T cells (*p* < 0.01), and reduced the percentage of B-cells (*p* < 0.05) resulting in an altered T-cell/B-cell ratio (**Figure 5B**). However, this effect was not apparent in infected pigs fed the BBE probiotics, with the T-cell/B-cell ratio equivalent to uninfected pigs, indicating that *O. dentatum*-induced alterations in lymphocyte populations were attenuated in these animals (**Figure 5B**). In contrast, LB probiotic supplementation did not have this modulatory effect, with no significant interaction and only a main effect of infection in analysis of both T-cell and B-cell populations (**Figure 5B**). Analysis of other cell populations, namely CD3^+^CD4^+^ helper and CD3^+^CD8^+^ cytotoxic T cells, or monocytes, showed no significant effects of either diet or infection (data not shown).

To assess functional cellular immune responses in peripheral and lymphoid tissues, PBMCs and CLN cells were stimulated with LPS or PHA, respectively, and cytokine secretion quantified. Infection did not consistently change the cytokine secretion pattern (**Figure 6**). In contrast, BBE supplementation substantially modulated cytokine profiles, although the effect was dependent on infection status. There was an interaction (*p* < 0.05) between probiotics and infection on mitogen-induced TNFα secretion from CLN cells, with BBE supplementation significantly reducing TNFα production in uninfected pigs, but not in infected animals. In contrast, IL-10 production tended to be enhanced by BBE in both infected and uninfected pigs (*p* = 0.06 for main effect of probiotic supplementation; **Figure 6A**). In PBMCs, BBE significantly suppressed LPS-induced IL-1β in both uninfected and infected pigs (*p* < 0.05; **Figure 6B**), with a similar tendency for IL-10 secretion (*p* = 0.06; **Figure 6B**). TNFα followed the same pattern but the differences were not significant (**Figure 6B**). There was an interaction (*p* < 0.05) between probiotics and infection for IL-6 production, with secretion reduced in uninfected pigs fed with BBE, but tended to be enhanced in infected pigs (**Figure 6B**). The effects of LB probiotics were less apparent. LB supplementation resulted in lower (*p* < 0.05) TNFα secretion from CLN cells, independently of infection status, but there were no effects on the other cytokines measured in either CLN or PBMC (**Supplementary Figure 3**). Collectively, these data suggest that BBE probiotics have an anti-inflammatory effect in the absence of parasite infection. However this effect was modulated in infected pigs. Whereas IL-1β was strongly suppressed in PBMC from both uninfected and infected animals receiving BBE, the effect on other cytokines such as IL-6 appeared to be influenced by the parasitic infection, with the suppressive effect less evident in infected pigs. These data suggest that concurrent helminth infection may restrict the anti-inflammatory properties of BBE probiotics.

**Figure 6.**
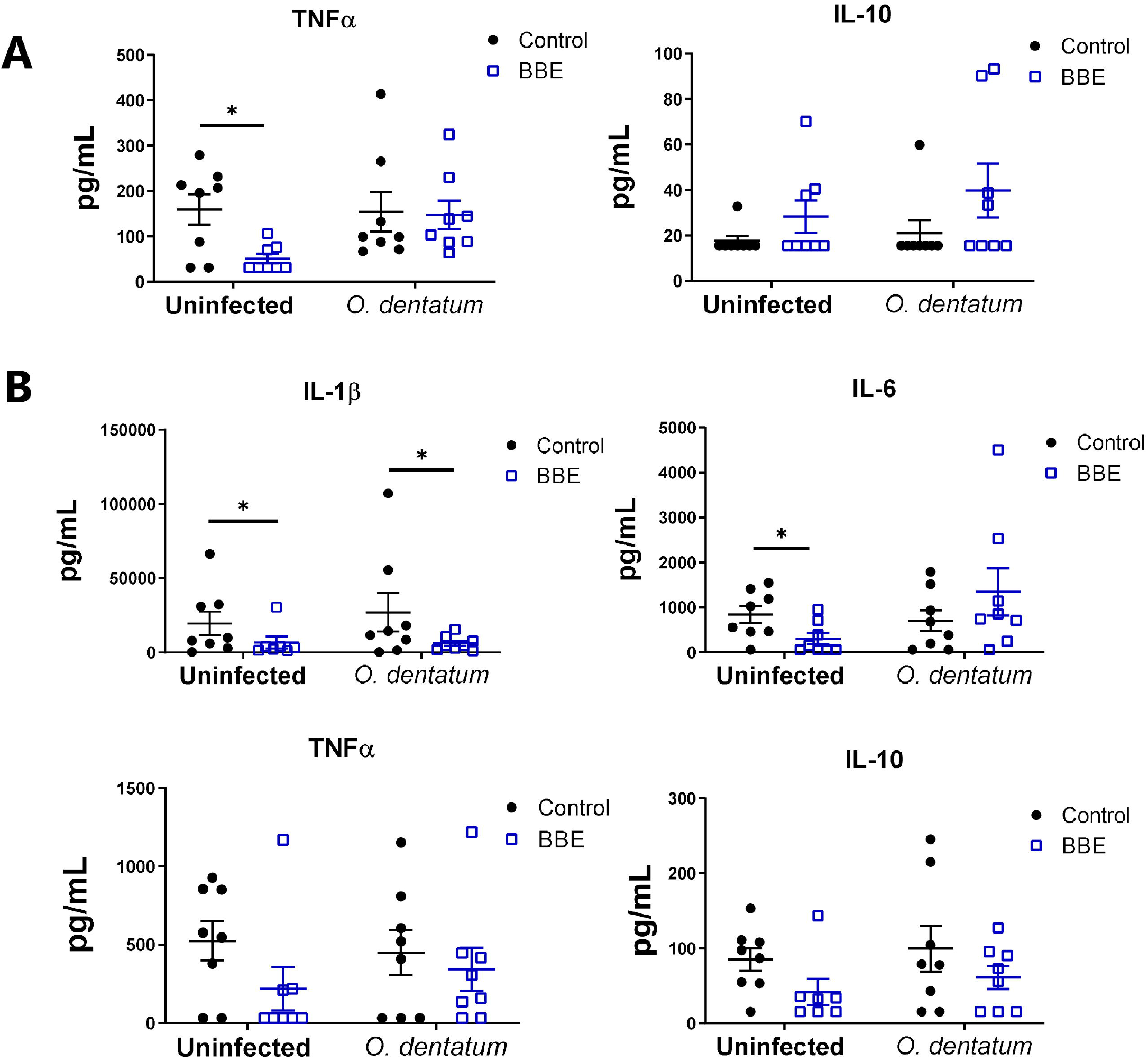
*Ex vivo* cytokine secretion is modulated by probiotics and *Oesophagostomum dentatum* infection. **A)** Phytohaemagglutinin-induced secretion of TNFα and IL-10 in ileal-caecal lymph node cultures. Pigs were either uninfected or infected with *O. dentatum* for 28 days, with or without supplementation of a mixture of *Enterococcus faecium* and *Bacillus* sp. (BBE). **B)** LPS-induced secretion of IL-1β, IL-6, TNF α and IL-10 in peripheral blood mononuclear cells from pigs infected with *O. dentatum* for 28 days or uninfected pigs, with or without supplementation of BBE. * *p* < 0.05 by GLM analysis. n=8 pigs per treatment group.

### Probiotics attenuate *O. dentatum-induced* inflammatory gene expression in the proximal colon

To explore in more detail if the dietary probiotics modulated local host immune responses, we investigated changes in gene expression in the proximal colon during *O. dentatum* infection. A panel of genes was selected to represent Th1-, Th2- and regulatory immune responses, as well as mucosal barrier and innate immunity-related genes. Principal component analysis (PCA) of the relative expression of all genes analysed in the proximal colon illustrated a marked effect of *O. dentatum* infection (**Figure 7A**), and a lesser influence of probiotic supplementation (**Figure 7B**). In the absence of probiotic supplementation, there was a prototypical type-2 polarised immune gene expression profile in the proximal colon of pigs infected with *O. dentatum*, relative to uninfected animals. Infection with *O. dentatum* significantly increased expression of *IL4, IL13, ARG1, CCL17* and *CCL26*, with a concurrent trend for down-regulation of the expression of Th1-related genes such as *IL8* (**Figure 7C**; **Supplementary Table 4**). In addition, increased expression of mucosal barrier-related genes, such as *RETNLB, FFAR2*, and *DCLK1*, and innate immune genes such as *IL6*, *C3* and *PTGS2* (encoding cyclooxygenase-2) were also observed in infected, control-fed pigs (**Figure 7C**; **Supplementary Table 4**).

**Figure 7.**
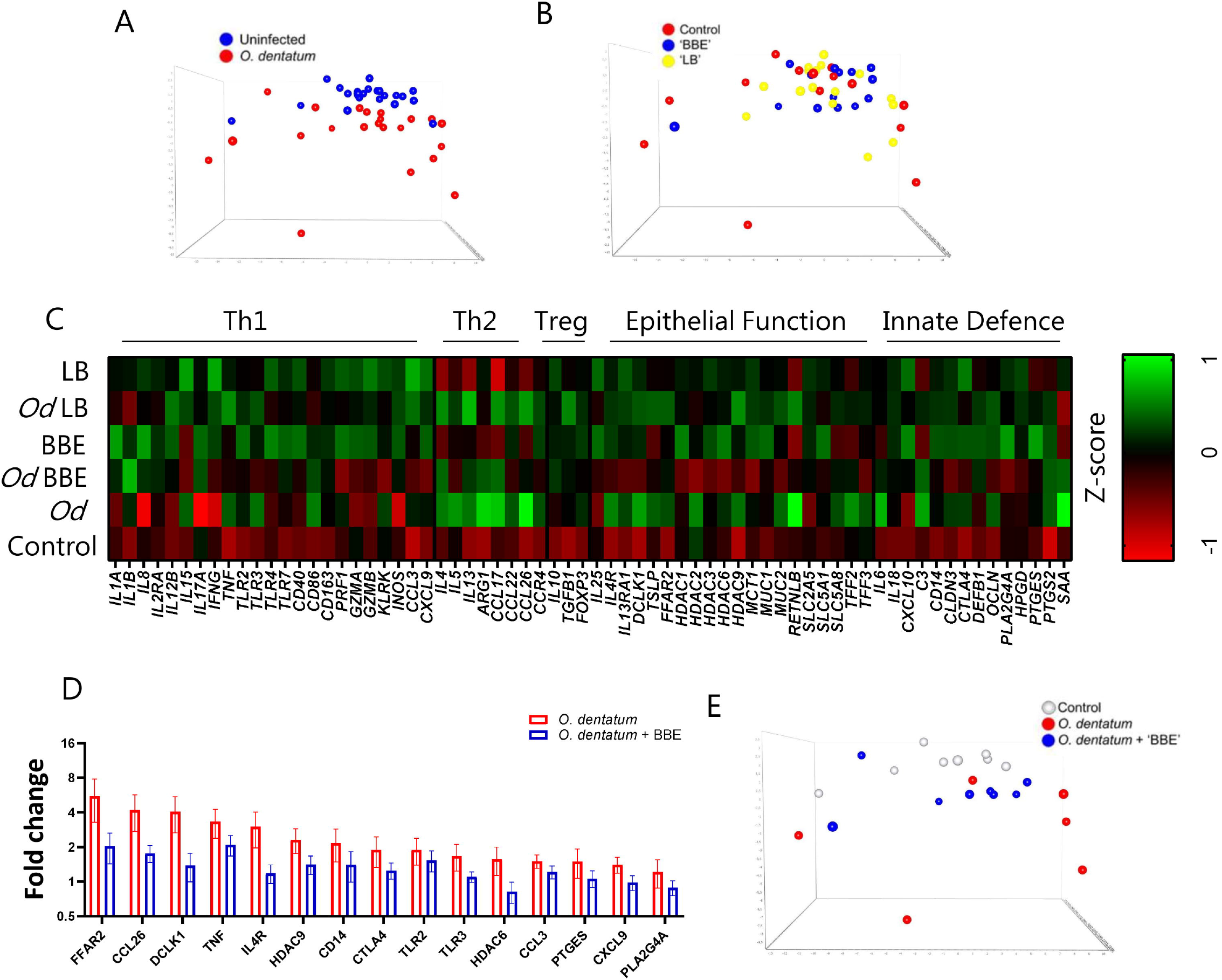
Probiotics and *Oesophagostomum dentatum* infection alters immune gene expression profiles. **A-B)** Principal component analysis of immune gene expression in the proximal colon at day 28 post-infection as a result of *O. dentatum* infection **(A)** or diet supplementation with probiotic mixtures *Enterococcus faecium* and *Bacillus* sp. (BBE). or LGG and Bb12 (LB) **(B)**. **C)** Expression of genes involved in different biological function as a result of *O. dentatum* infection (Od), BBE or LB supplementation, or *O. dentatum* infection combined with BBE or LB supplementation. The control group received no infection or probiotic treatment. Data presented as Z-scores of relative gene expression data. **D)** Fold changes in expression of genes from proximal colon tissue significantly altered (*p* < 0.05) by the interaction of *Oesophagostomum dentatum* infection and dietary supplementation with a mixture of *Enterococcus faecium* and *Bacillus* sp. (BBE), in comparison to control-fed, *O. dentatum*-infected controls. n=8 pigs per treatment group. **E)** Principal component analysis showing immune gene expression in the proximal colon at day 28 post-infection in control pigs (no infection or probiotics), *O. dentatum* infection without probiotics, and *O. dentatum* with BBE supplementation.

We noted a moderately enhanced Th1 polarization as a result of probiotic supplementation. Both probiotic treatments increased the expression of *IL8*, *IL12B* and *INOS* in both uninfected and *O. dentatum*-infected animals. LB supplementation also significantly increased *IFNG* expression (**Figure 7C**; **Supplementary Table 4**), as well as *CXCL10* expression but only in males (*p* < 0.05 for interaction between sex and LB supplementation).

Strikingly, in *O. dentatum-infected* pigs, BBE supplementation markedly attenuated the helminth-induced increases in gene expression relative to control-fed animals In BBE-fed pigs, Th2 genes were still up-regulated as a result of helminth infection, but to a lesser degree compared to *O. dentatum* infected pigs fed only the control diet (**Figure 7C**). For genes where there was a significant interaction (*p* < 0.05) between BBE supplementation and infection, in every case this resulted in significant down-regulation of expression in infected, BBE-fed pigs compared to infected, control-fed pigs (**Supplementary Table 4**). This included key Th2 and epithelial/ mucosal barrier related genes, including those coding for the short-chain fatty acid receptor *FFAR2*, the epithelial cell kinase and tuft cell marker *DCLK1* the interleukin-4 receptor *IL4*, and the eosinophil chemoattractant *CCL26* (**Figure 7D**). Moreover, the helminth-induced expression of other immune related genes such as *TNF, CTLA4* and *PLA2G4A* was significantly attenuated by BBE supplementation (**Figure 7D**). This was evident in PCA analysis which showed that *O. dentatum-infected* pigs administered BBE clustered closer to uninfected control pigs than *O. dentatum*-infected pigs without probiotic supplementation, suggesting that the response to infection was muted in these animals, and that BBE acted to restrain the localized inflammatory response to the parasite (**Figure 7E**). A similar pattern was evident in infected pigs with LB supplementation, but the effect was less pronounced, with the immune gene expression profiles with the immune gene profile more closely resembling that of *O. dentatum*-infected pigs fed the control diet (**Figure 7E**). However, we did note a trend (*p* < 0.1) for interactions between infection and LB supplementation for the expression, of *ARG1, TLR3, IL1B*, and *CTLA4*, with the infection-induced expression of these genes being attenuated to some extent by LB (**Figure 7C; Supplementary Table 4)**. Thus, probiotic supplementation (most primarily with BBE) acted to attenuate parasite-induced, type-2 biased inflammatory responses in the colon.

## Discussion

The beneficial effect of probiotics on health and control of bacterial infections is well-documented, however the potential interactions of probiotics with helminth infection and the mechanisms by which they can influence mucosal immune responses is not well understood. We found that probiotics (BBE in particular) were capable of supressing *ex vivo* inflammatory cytokine production and attenuating the host mucosal immune responses elicited in response to infection. Neither probiotic mixture modulated the establishment or infection kinetics of *O. dentatum*. However, both mixtures appeared to beneficially modulate the intestinal microbiota composition, as evidenced by increased bacterial diversity in both faecal and large intestinal samples. Interestingly, we noted that these effects were to some extent modulated by *O. dentatum* infection, suggesting a novel interaction of parasite infection on probiotic activity. Furthermore, we observed attenuation of the prototypical type-2 inflammation induced by *O. dentatum* by BBE probiotics.

*O. dentatum* infection is highly prevalent in pigs worldwide. Whereas dietary prebiotics, such as inulin, have been shown to be highly effective in reducing parasite burdens, our results here show that supplementation of these specific probiotic strains did not have an anti-parasitic effect. The mode-of-action of prebiotics against *O. dentatum* is hypothesized to result from a selective enrichment of lactic acid producing bacteria, and production of GM-derived metabolites such as SCFA, which lower the colon pH and create an inhospitable environment for helminths ^26^. Despite an increase in D-lactic acid induced by LB, we did not observe changes in gut pH (or total SCFA levels) as a result of either probiotic mixture. Thus, the administration of certain probiotic bacteria was insufficient to have an anthelmintic effect, although associated effects on the immune system or GM may still markedly impact gut health.

Helminth infection is typically associated with a rise in antibody secretion and the initiation of a characteristic Th2 immune response. Similarly to Andreasen *et al*. (2015) ^27^ we observed a type-2 immune response in control-fed pigs infected with *O. dentatum*, with increased antibody secretion, peripheral T cell activation, and type-2 immune gene expression profiles in the proximal colon confirming an active host immune response was elicited. Interestingly, infected pigs fed BBE probiotics exhibited a reduction in epithelial immune genes, such as *TSLP, IL4R* and *FFAR2*, compared to the *O. dentatum*-infected pigs fed only the control diet. In addition, BBE treatment alone tended to reduce expression of key Th2 immune genes, such as *IL4*, *IL5* and *CCL26*, and appeared to diminish the parasite-induced increase in the expression of these genes in infected pigs fed BBE. Together, this suggests that the typical polarised helminth-mediated Th2 immune response is attenuated by the supplementation of *Bacillus* spp. plus *E. faecium-based* probiotics. This attenuation of prototypical helminth-induced immune response has been observed previously in *A. suum*-infected pigs fed *L. rhamnosus* LGG ^22^. Jang *et al*. (2017) reported reduced IgG2 antibody titres and reduced expression of *IL13*, eosinophil peroxidase *EPX*, and *CCL26* in *A. suum*-infected pigs supplemented with LGG. The observed suppression of Th2 and epithelial gene expression profiles in this study may have been the result of the probiotics exerting a regulatory effect to maintain intestinal immune homeostasis.

We observed that probiotic supplementation appeared to significantly alter the intestinal microbiota, with both mixtures (BBE and LB) improving the microbial diversity and richness over the course (day 28 post-infected compared to 7 days pre-infection) of the study and at different segments of the intestinal tract. PERMANOVA analysis confirmed that probiotic supplementation did have a modulatory effect on the microbiota, although the changes could not be ascribed to specific taxa. The modest impact of probiotics on the composition of the GM appears to be in keeping with several studies that reported minor compositional alterations as a result of supplementation with a range of probiotic strains ^6, 28^. Interestingly, both probiotic mixtures induced subtle alterations to SCFA and lactic acid levels present in intestinal digesta, suggesting that even with limited changes in the GM, potentially beneficial outcomes to intestinal health can still be achieved, as was evident by the modulation of intestinal immune gene expression profiles.

To our knowledge this is the first time the porcine GM has been characterised during *O. dentatum* infection. Consistent with previous observations in pigs infected with *T. suis* ^19, 20^, *O. dentatum* infection altered β-diversity in the caecum and colon. However, unlike *T. suis*, this modulation did not appear to be associated with defined bacterial taxa, and significant changes were not observed in faeces or the small intestine. This suggests that *O. dentatum* infection had a localised impact on the GM without inducing changes throughout the intestinal tract. The most striking observation was the apparent ability of *O. dentatum* to suppress the changes in the GM brought about by probiotics that were observed in uninfected pigs. Thus, concurrent parasitic infections, which are common in livestock and humans in developing countries, may be a previously unappreciated factor influencing the health benefits of dietary probiotics.

The mechanisms by which probiotics alter the response to helminth infection requires further investigation. Various modes-of-action have been proposed for the health benefits of probiotic bacteria. Probiotics may adhere to intestinal epithelial cells and thereby prevent the attachment of potentially pathogenic bacteria such as *E. coli*, as well as inducing mucus production and the stimulation of antimicrobial peptides ^29^. Furthermore, probiotics may regulate inflammatory responses by binding to PRRs on immune cells and promoting secretion of IL-10 or TGF-β, which can supress inflammatory cytokine production ^30^. Moreover, probiotics such as LGG have previously been shown to promote Th1 responses in pigs, and the Th1-stimulating properties of probiotics has been suggested to underlie the ability of probiotics to supress symptoms of allergies in humans and animal models ^30, 31^. Indeed, our gene expression data in the colon indicated a modest Th1-polarizing effect of both probiotic mixtures in the absence of infection, suggesting that host pattern recognition receptors recognize the bacteria and respond with production of type-1 cytokines and innate immune mediators that are typically produced in response to TLR or NOD receptor binding. Probiotics have also been shown to induce regulatory responses that can alleviate inflammation during pathogen challenge in pigs ^32^, and thus the attenuation of the helminth-induced type-2 response may then derive from the ability of the probiotic bacteria to restore homeostasis in the face of acute pathogen-driven inflammation. Interestingly, we observed that BBE appeared to be more efficient than LB in modulating host immune responses, which may reflect the inclusion of porcine-derived strains in the BBE mixture.

In conclusion, we show here that probiotics, in particular the strains *Bacillus amyloliquefaciens, B. subtilis*, and *Enterococcus faecium*, do not appear to directly affect worm establishment and development but do regulate inflammatory responses and attenuate host mucosal immune function during *O. dentatum* infection, which may serve to regulate host intestinal function and maintain immune homeostasis. This probiotic-mediated regulation of host immune responses is also indicative of the ability of probiotics to potentially dampen Th2-mediated pathology as a result of, for example, food allergies ^33–35^. Moreover, the ability of these probiotic strains to attenuate pathogen-induced inflammatory responses may have relevance for dietary interventions that seek to maintain intestinal homeostasis during infectious challenge.

## Materials and methods

### Experimental design

A total of 48 Yorkshire-Landrace pigs (females and castrated males, 8-10 weeks old, initial body weight approximately 20 kg) were sourced from a specific pathogen-free farm. After stratification based on sex and weight, pigs were randomly allocated to one of six groups. Each treatment group contained eight pigs housed in a separate pen. Two groups (each n=8) received the basal diet only (based on ground barley and soybean containing 16.2% crude protein). Two groups (each n=8) received the same diet supplemented with BBE containing the strains *Bacillus amyloliquefaciens* 516 (porcine origin)*, B. subtilis* 541 (human origin), and *Enterococcus faecium* 669 (human origin). The final two groups (each n=8) received the basal diet supplemented with LB, containing the strains *Lactobacillus rhamnosus* LGG^®^ (human origin; DSM33156) and *Bifidobacterium animalis* subsp. *Lactis* BB-12^®^ (food origin; DSM15954). All probiotic strains were supplied by Chr.Hansen A/S, Denmark. Pigs were fed twice a day with the probiotic-supplements mixed with the standard feed immediately before feeding. For both probiotic mixtures, pigs received 2 x 10^10^ CFU per day.

After two weeks of diet adaptation, a total of 24 pigs (8 pigs from each diet treatment group) were each inoculated with 25 *O. dentatum* third stage larvae (L3)/ kg body weight, by oral gavage. These pigs subsequently continued to receive the same *O. dentatum* L3 dose three days a week until study end (a total of four weeks). Infection doses were provided during the morning feeding, and were uniformly distributed on top of the feed. The dosed feed was provided in troughs that allowed all pigs’ adequate space to feed equally and simultaneously. The dosing regime was chosen to mimic a natural moderate exposure level and the average approximate theoretical total dose during the study was 22,000 *O. dentatum* L3/ pig. The remaining 24 pigs were uninfected for the duration of the study.

All pigs had been vaccinated against *L. intercellularis* with one dose of a live, attenuated vaccine (Enterisol^®^ Ileitis, Boehringer Ingelheim) on farm four weeks prior to arriving on the experimental premises (which was six weeks prior to infection with *O. dentatum*). All pigs were confirmed negative for *O. dentatum* infection upon arrival by McMaster faecal egg count and serology. For the duration of the study, all pigs were housed on concrete floored pens with wood chips and water provided *ad libitum*. Welfare checks were performed daily, with body weight monitored and reported weekly. At day 28 post-infection (p.i.), 48 pigs were sacrificed over the course of three days by stunning with captive bolt followed by exsanguination.

This study was approved by the Danish Animal Experimentation Inspectorate (License number 2015-15-0201-00760), and performed at the Experimental Animal Unit, University of Copenhagen according to FELASA guidelines and recommendations.

### Digesta sampling and *O. dentatum* isolation

Weekly blood and faecal samples were taken between arrival (day −14) and until the end of the study (day 28 p.i.). Blood samples were taken in order to collect serum for ELISA (see below), and isolate peripheral blood mononuclear cells (PBMCs; day 28 p.i. only). Faecal samples were scored following a 5-point scale (1 – hard; 2 – normal; 3 – soft; 4 – watery; 5 – diarrhoea) in order to monitor changes in faecal consistency as a result of probiotic supplementation. After scoring, samples were cooled to ~4°C immediately upon collection for subsequent enumeration of *O. dentatum* egg counts per gram of faeces (EPG) using a McMaster faecal egg count method (as described in Roepstorff & Nansen, 1998) ^36^.

At necropsy, fresh intestinal digesta samples were collected from specific intestinal sections: jejunum (mid-point of the small intestine), ileum (10 cm proximal from ileocaecal junction), caecum, proximal colon (20 cm distal from ileocaecal junction) and distal colon (central part of the spiral) colon) for microbiota and pH measurement, with additional samples taken from the proximal colon for SCFA analysis, as previously described ^19^. Following this, *O. dentatum* larvae and adults were recovered according to the agar-gel migration technique described previously by Slotved *et al*. (1996) ^37^. Briefly, luminal contents of caecum and colon were collected and diluted to a total volume of 10 litres using 0.9% saline (37°C). A 5% sub-sample was then embedded in 2% agar on cloths that were then suspended in saline and incubated for 24 hours at 37°C to isolate immature and adult *O. dentatum* from each pig. Worms were isolated on a 38 μm mesh and stored in 70% ethanol for later enumeration. For each pig, ten adult female and male worms were selected for length measurement, using Leica Application Suite version 4.7 (Leica Microsystems, Germany), as a measure of *O. dentatum* fitness.

### Cell isolation, flow cytometry and assessment of cytokine production

Ileo-caecal lymph nodes (CLNs) were dissected and passed through a 70 μM cell strainer to obtain single cell suspensions. After a series of washing, the cells were prepared for flow cytometric phenotypic analysis of T cells, B cells and monocyte populations as described in Myhill *et al*. (2018) [15]. Flow cytometry was performed using a BD Accuri C6 flow cytometer (BD Biosciences), and data were analysed using Accuri CFlow Plus software (Accuri^®^ Cytometers Inc., MI, USA). PBMCs were isolated from heparinised whole blood using Histopaque-1077 (Sigma-Aldrich) and centrifugation. To assess cytokine production, isolated CLN cells were cultured for 48 hours in complete media (RPMI 1640 supplemented with 2 mM L-glutamine, 10% calf serum, 100μg/mL streptomycin and 100 U/mL penicillin) together with 10 μg/mL phytohemagglutinin (Sigma-Aldrich). Measurement of secreted TNFα and IL-10 was assessed using commercial ELISA kits (R&D systems). Isolated PBMCs in complete media were stimulated with LPS (1 μg/mL), cultured for 24 hours, and concentrations of IL-6, TNFα, IL-10 and IL-1β assessed by ELISA. Values below the detection limit were assigned an arbitrary value of half the lowest value of the standard curve.

### *O. dentatum* culture

*O. dentatum* larvae were isolated from infected control-fed pigs, and washed extensively in 37°C saline. The exsheathed larvae were cultured in complete media containing antibiotics and fungicide for 3 days at 37°C to obtain excretory/secretory (E/S) products. Every day the culture media was removed and stored at −80°C, and replaced with fresh media. Pooled culture media containing E/S was concentrated by centrifugation using Amicon ultra centrifugal filter units (MWCO 10 kDa, Sigma-Aldrich, Denmark), and filtered prior to testing of protein content by bicinchoninic (BCA) assay (Thermo Fisher Scientific).

### *O. dentatum* ELISA

Anti-*O. dentatum* IgA and IgG_1_ levels in serum were quantified by ELISA as described in Myhill *et al*. (2018) ^19^. Briefly, plates (Nunc Maxisorb) were coated with 5 μg/mL *O. dentatum* larval E/S overnight at 4°C. Serum antibodies were then detected using goat anti-pig IgA-horseradish peroxidase (HRP; BioRad, Germany), or mouse anti-pig IgG_1_ (clone K139-3C8; BioRad) followed by goat anti-mouse IgG-HRP conjugate (BioRad). Incubations were for 1 hour at 37°C, and between all steps, plates were washed four times with PBS plus 0.02% Tween 20. After development with tetramethylbenzidine (TMB) substrate, the reaction was stopped with 0.2M H_2_SO_4_, and the plates read at 450 nM with a Multiskan FC plate reader (Waltham, Massachusetts, USA).

### Quantitative real-time PCR

Total RNA was extracted from proximal colon tissue using a miRNAeasy^®^ Mini kit (Qiagen, CA, USA) according to manufacturer’s guidelines, and as described in Myhill *et al*. (2018) ^19^. Synthesis of cDNA and pre-amplification was conducted as described in Williams *et al*. (2017) ^24^. A panel of 77 genes of interest, including key Th1/Th2/Treg/innate immune response-related genes and epithelial/mucosal barrier function-related genes, were examined on a BioMark HD Reader (Fluidigm). First, a thermal mix and hot start protocol was performed to mix primers, samples and reagents (50°C for 2 min, 70°C for 30 min, 25°C for 10 min, 50°C for 2 minutes, 95°C for 10 min), followed by qPCR using the following cycling conditions of: 35 cycles at 95°C for 15 seconds and 60°C for 1 min. After data pre-processing, 68 genes of interest passed quality control criteria and were statistically analysed. Normalization using several validated reference housekeeping genes and data pre-processing, was carried out as described in Skovgaard *et al*. (2009) ^38^. Primer sequences are presented in **Supplementary Table 5.**

### 16S rRNA sequencing of microbiota

DNA was extracted from faeces or intestinal content in a randomized order using the Bead-Beat Micro AX Gravity Kit (A&A Biotechnology, Poland) according to manufacturer’s instructions. Prior to extraction, samples were lysed in LSU buffer supplemented with Lysozyme (4000 U) and Mutanolysin (50 U), and incubated at 50°C for 20 min. The concentration and purity of extracted DNA were determined using a NanoDrop ND-1000 spectrophotometer and normalized to 10 ng/μl. High throughput sequencing based 16S rRNA gene amplicon (V3-region) sequencing was carried out on an Illumina NextSeq platform as previously described ^39^.

The raw dataset containing pair-ended reads with corresponding quality scores were merged and trimmed using fastq_mergepairs and fastq_filter scripts implemented in the USEARCH pipeline as described previously ^39^. Purging the dataset from chimeric reads and constructing zero radius Operational Taxonomic Units (zOTU) was conducted using UNOISE. The Greengenes (13.8) 16S rRNA gene collection was used as a reference database. Quantitative Insight Into Microbial Ecology (QIIME) open source software package (v2019.7.0) was used for subsequent analysis steps ^40^. Alpha diversity measures: observed species (number of zOTUs) and Shannon diversity indices were computed for rarefied OTU tables (10,000 reads/sample) using the alpha rarefaction workflow. Differences in alpha diversity were determined using a t-test-based approach employing the non-parametric (Monte Carlo) method (999 permutations) implemented in the compare alpha diversity workflow. Principal Coordinates Analysis (PCoA) plots were generated with the Jackknifed Beta Diversity workflow based on 10 distance metrics calculated using 10 sub-sampled OTU tables. The number of sequences taken for each jackknifed subset was set to 85% of the sequence number within the most indigent sample (~ 10,000). Community differences (beta-diversity) were revealed by weighted and unweighted Unifrac distance metrics visualised as Principle Coordinate Analysis (PCoA) plots. Permutational Multivariate Analysis of Variance (PERMANOVA) and Non-parametric microbial interdependence test (NMIT) were used to evaluate group differences based on weighted and unweighted UniFrac distance matrices. Taxa-level differences were assessed using longitudinal feature-volatility analysis and analysis of composition of microbes (ANCOM).

### Statistical analysis

Data were analysed using general linear model (GLM) using IBM SPSS Statistics 28. For each separate probiotic mixture (BBE or LB), the effects of probiotic supplementation and parasite infection, and their interaction, were compared to control-fed animals using a separate factorial analysis. The model included infection status, probiotic supplementation and sex as fixed factors, together with their first-order interactions. Sex was removed from the model when not significant. For analysis of ELISA data, time was included as an additional fixed factor to account for repeated measurements. Assumptions of normality were checked through inspection of histogram plots and Shapiro-Wilk and Kolmogorov-Smirnov tests of GLM residuals, and data that did not conform to normality was transformed with either square-root or log_10_ transformations prior to analysis. Significance was taken at *p* < 0.05, and a trend at *p* <0.1.

## Acknowledgements

We thank L. Christensen (Department of Veterinary and Animal Sciences, University of Copenhagen) for her practical support throughout the study. We also thank K. Tarp (Department of Biotechnology and Biomedicine, Technical University of Denmark) for technical assistance with qPCR. In addition, we would like to thank Bea Nielsen (Chr. Hansen A/S) for providing all probiotic strains/ mixtures used within this study, and for technical advice.

## Data Availability Statement

Raw sequence data is available at Sequence Read Archive (https://www.ncbi.nlm.nih.gov/sra/) under the accession number PRJNA746763. All other data is available within the manuscript or supplementary material.

