## Supplementary Information for "Parasite-probiotic interactions in the gut: *Bacillus* sp. and *Enterococcus faecium* regulate type-2 inflammatory responses and modify the gut microbiota of pigs during helminth infection"

**Contents:**

**Supplementary Figures 1-3**

**Supplementary Tables 1-5**

*A*

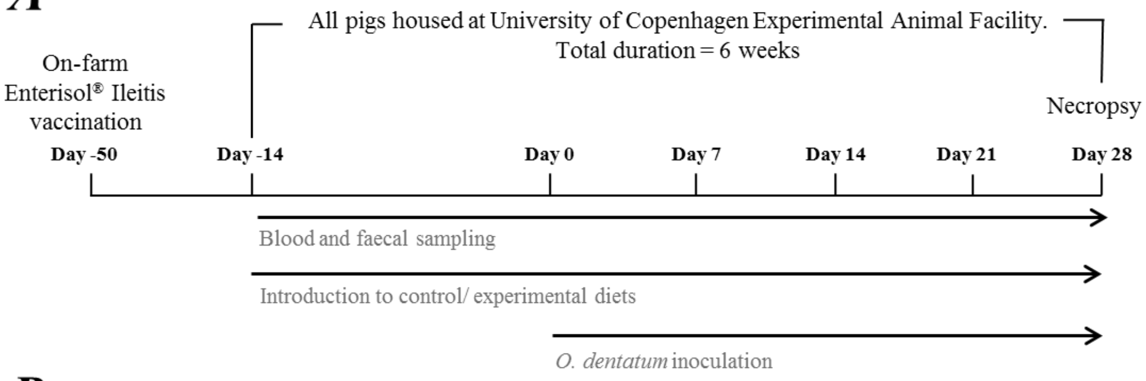


*B*

| | **Standard, non-pelleted feed ('Control')** | **Standard feed + Probiotic Mix 1 ('BBE')** | **Standard feed + Probiotic Mix 2 ('LB')** |
|---|---|---|---|
| **No infection** | n = 8 | n = 8 | n = 8 |
| ***Oesophagostomum dentatum* infection** | n = 8 | n = 8 | n = 8 |

**Supplementary Figure 1.** Experimental set up. (A) At day -14, 48 pigs arrived and were fed one of three diets. At day 0, 24 pigs were inoculated with 25 *O. dentatum* third stage larvae (L3) / kg body weight, followed by similar inoculations three times a week until day 28 post-infection (p.i.). (B) Final number of animals per treatment group at termination of study at day 28 p.i.

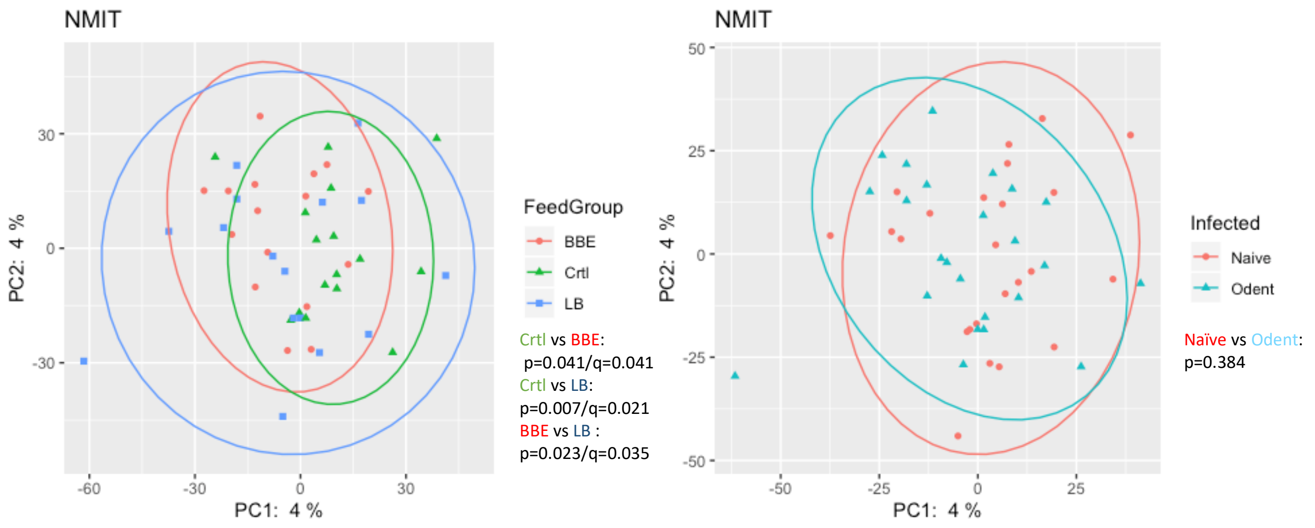


**Supplementary Figure 2**

*Figure S1:* Pooled NMIT analysis according to probiotic supplementation (left) or *Oesophagostomum dentatum* (*Odent*) (right).

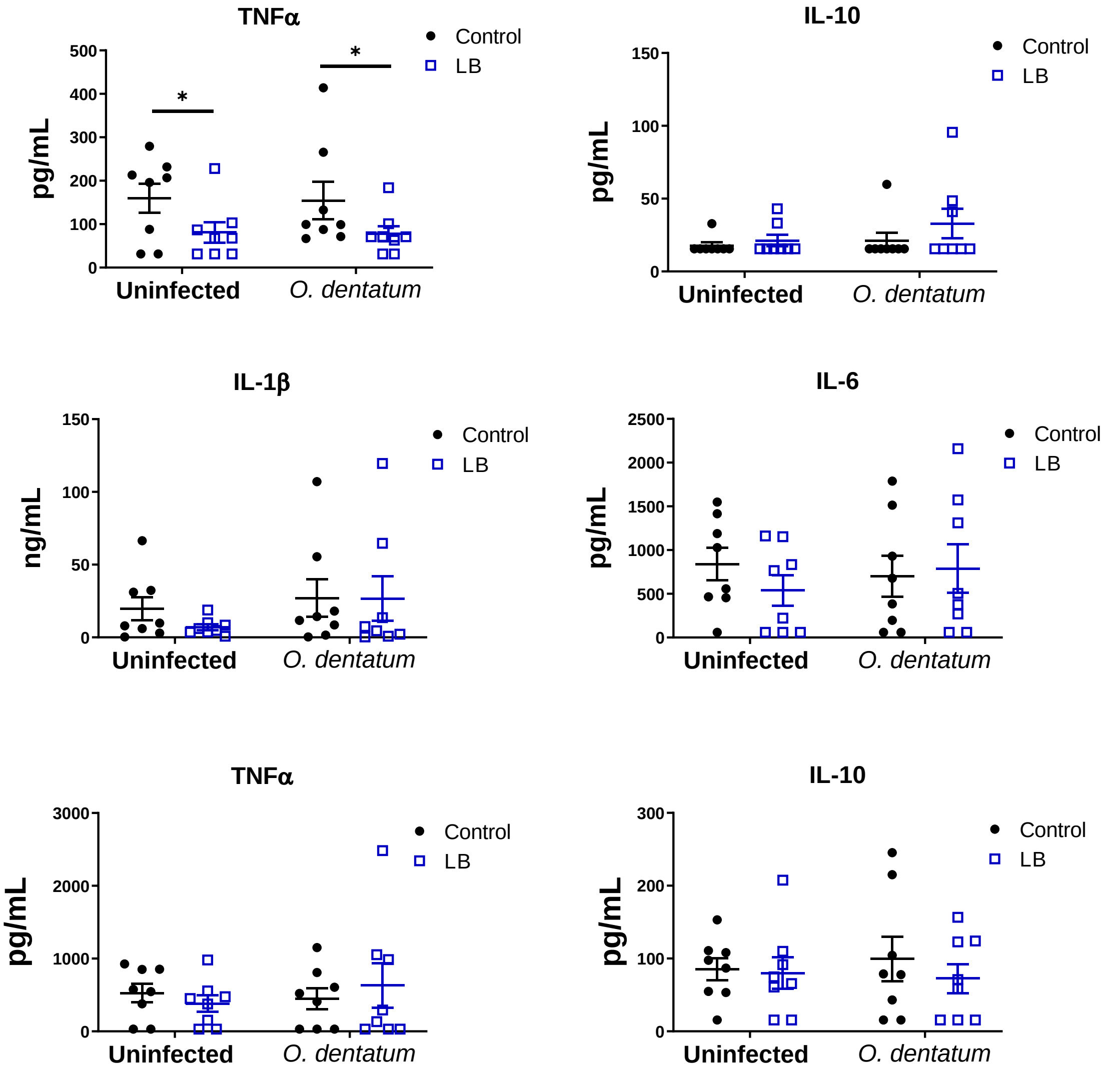


**Supplementary Figure 3.** ***Ex vivo*** **cytokine secretion in pigs given LGG/Bb12**

**A)** Phytohaemagglutinin-induced secretion of TNFα and IL-10 in ileal-caecal lymph node cultures. Pigs were either uninfected or infected with *O. dentatum* for 28 days, with or without supplementation of a mixture of LGG and Bb12 (LB). **B)** LPS-induced secretion of IL-1β, IL-6, TNF α and IL-10 in peripheral blood mononuclear cells from pigs infected with *O. dentatum* for 28 days or uninfected pigs, with or without supplementation of LB. * $p < 0.05$ by GLM analysis. n=8 pigs per treatment group.

**Supplementary Table 1.**

Alpha diversity indices (Faiths PD) for BBE (left) and LB (right) groups for each segment. Pairwise Kruskal-Wallis. Dark grey: p > 0.1; Light grey: p 0.05 to 0.099; White: p< 0.05

| BBE | | |
|---|---|---|
| | | **Faiths PD** |
| **JEJUNUM** | Crtl vs Od | 0.834 |
| | Crtl vs BBE | 0.753 |
| | Crtl vs BBE+Od | 0.728 |
| | Od vs BBE | 0.834 |
| | Od vs BBE+Od | 0.165 |
| | BBE vs BBE+Od | 0.083 |
| **ILEUM** | Crtl vs Od | 0.668 |
| | Crtl vs BBE | 0.105 |
| | Crtl vs BBE+Od | 0.728 |
| | Od vs BBE | 0.197 |
| | Od vs BBE+Od | 0.519 |
| | BBE vs BBE+Od | 0.036 |
| **CAECUM** | Crtl vs Od | 0.439 |
| | Crtl vs BBE | 0.156 |
| | Crtl vs BBE+Od | 0.317 |
| | Od vs BBE | 0.529 |
| | Od vs BBE+Od | 0.817 |
| | BBE vs BBE+Od | 0.355 |
| **PROX** | Crtl vs Od | 0.908 |
| | Crtl vs BBE | 0.345 |
| | Crtl vs BBE+Od | 0.355 |
| | Od vs BBE | 0.563 |
| | Od vs BBE+Od | 0.406 |
| | BBE vs BBE+Od | 0.908 |
| **DISTAL** | Crtl vs Od | 0.046 |
| | Crtl vs BBE | 0.009 |
| | Crtl vs BBE+Od | 0.023 |
| | Od vs BBE | 0.208 |
| | Od vs BBE+Od | 0.752 |
| | BBE vs BBE+Od | 0.115 |

| LB | | |
|---|---|---|
| | | **Faiths PD** |
| **JEJUNUM** | Crtl vs Od | 0.916 |
| | Crtl vs LB | 0.208 |
| | Crtl vs LB+Od | 0.487 |
| | Od vs LB | 0.059 |
| | Od vs LB+Od | 0.298 |
| | LB vs LB+Od | 0.643 |
| **ILEUM** | Crtl vs Od | 0.568 |
| | Crtl vs LB | 0.654 |
| | Crtl vs LB+Od | 0.563 |
| | Od vs LB | 0.568 |
| | Od vs LB+Od | 0.606 |
| | LB vs LB+Od | 0.728 |
| **CAECUM** | Crtl vs Od | 0.606 |
| | Crtl vs LB | 0.199 |
| | Crtl vs LB+Od | 0.156 |
| | Od vs LB | 0.418 |
| | Od vs LB+Od | 0.401 |
| | LB vs LB+Od | 0.817 |
| **PROX** | Crtl vs Od | 0.728 |
| | Crtl vs LB | 0.172 |
| | Crtl vs LB+Od | 0.916 |
| | Od vs LB | 0.418 |
| | Od vs LB+Od | 0.728 |
| | LB vs LB+Od | 0.401 |
| **DISTAL** | Crtl vs Od | 0.046 |
| | Crtl vs LB | 0.005 |
| | Crtl vs LB+Od | 0.093 |
| | Od vs LB | 0.366 |
| | Od vs LB+Od | 0.834 |
| | LB vs LB+Od | 0.699 |

**Supplementary Table 2.**

Beta diversity (unweighted UniFrac) for BBE groups for each segment. Permanova (pairwise Kruskal-Wallis) for DMs in Figure 3. Dark grey: p > 0.1; Light grey: p 0.05 to 0.099; White: p< 0.05

| BBE | | | |
|---|---|---|---|
| | | Unweighted | |
| | | p-value | q-value |
| JEJUNUM | Crtl vs BBE | 0.290 | 0.290 |
| | Crtl vs BBE+Od | 0.080 | 0.134 |
| | BBE vs BBE+Od | 0.089 | 0.134 |
| ILEUM | Crtl vs BBE | 0.026 | 0.039 |
| | Crtl vs BBE+Od | 0.163 | 0.163 |
| | BBE vs BBE+Od | 0.026 | 0.039 |
| CAECUM | Crtl vs BBE | 0.032 | 0.096 |
| | Crtl vs BBE+Od | 0.103 | 0.155 |
| | BBE vs BBE+Od | 0.208 | 0.208 |
| PROXIMAL | Crtl vs BBE | 0.003 | 0.006 |
| | Crtl vs BBE+Od | 0.026 | 0.026 |
| | BBE vs BBE+Od | 0.004 | 0.006 |
| DISTAL | Crtl vs BBE | 0.005 | 0.008 |
| | Crtl vs BBE+Od | 0.014 | 0.014 |
| | BBE vs BBE+Od | 0.005 | 0.008 |

**Supplementary Table 3.**

Beta diversity (unweighted UniFrac) for LB groups for each segment. Permanova (pairwise Kruskal-Wallis) for DMs in Figure 3. Dark grey: p > 0.1; Light grey: p 0.05 to 0.099; White: p< 0.05

| LB | | | |
|---|---|---|---|
| | | Unweighted | |
| | | p-value | q-value |
| JEJUNUM | Crtl vs LB | 0.067 | 0.099 |
| | Crtl vs LB+Od | 0.022 | 0.066 |
| | LB vs LB+Od | 0.099 | 0.099 |
| ILEUM | Crtl vs LB | 0.356 | 0.356 |
| | Crtl vs LB+Od | 0.073 | 0.110 |
| | LB vs LB+Od | 0.030 | 0.090 |
| CAECUM | Crtl vs LB | 0.049 | 0.074 |
| | Crtl vs LB+Od | 0.030 | 0.074 |
| | LB vs LB+Od | 0.276 | 0.276 |
| PROXIMAL | Crtl vs LB | 0.006 | 0.018 |
| | Crtl vs LB+Od | 0.095 | 0.095 |
| | LB vs LB+Od | 0.030 | 0.045 |
| DISTAL | Crtl vs LB | 0.008 | 0.014 |
| | Crtl vs LB+Od | 0.009 | 0.014 |
| | LB vs LB+Od | 0.018 | 0.018 |

**Supplementary Table 4. Relative expression and significance (*p*-value) of genes significantly influenced by diet, infection or interaction of both treatments. Significance determined as *p* ≤ 0.05. # indicates a trend of effect where *p* ≤ 0.1.**

Statistical analysis was conducted separately for each probiotic treatment, using a GLM analysis comparing the effect of probiotic supplementation and infection (and their interaction) to the control-diet groups (no probiotics).

| Immune function | Immune gene | Relative expression | | | | Significance (*p*-value) | | | | | Significance (*p*-value) | | |
|---|---|---|---|---|---|---|---|---|---|---|---|---|---|
| | | Control | *O. dentatum* | BBE | *O. dentatum* + BBE | Diet | Infection | Interaction | LB | *O. dentatum* LB | Diet | Infection | Interaction |
| Th1 | *IL1A* | 5.3 | 4.7 | 7.4 | 6.5 | 0.048 | | | 5.9 | 5.6 | | | |
| | *IL1B* | 2.2 | 3.2 | 3.3 | 4.4 | # 0.059 | # 0.078 | | 3.0 | 2.3 | | | 0.027 |
| | *IL8* | 7.7 | 5.5 | 11.0 | 8.7 | 0.007 | # 0.054 | | 9.4 | 7.9 | # 0.098 | # 0.082 | |
| | *IL12B* | 3.1 | 3.2 | 5.8 | 5.5 | 0.026 | | | 4.4 | 6.0 | 0.016 | | |
| | *IFNG* | 3.9 | 3.1 | 5.4 | 4.0 | | | | 6.4 | 5.2 | 0.002 | | |
| | *TNF* | 8.9 | 29.6 | 19.1 | 18.6 | | | 0.014 | 26.9 | 31.2 | # 0.091 | 0.026 | |
| | *TLR2* | 4.1 | 7.8 | 7.5 | 6.3 | | | 0.023 | 5.8 | 7.1 | | # 0.054 | |
| | *TLR3* | 1.8 | 3.0 | 2.3 | 2.0 | | | 0.047 | 2.2 | 2.6 | | 0.005 | # 0.054 |
| | *INOS* | 13.7 | 9.7 | 22.9 | 17.9 | # 0.076 | | | 19.1 | 21.4 | 0.034 | | |
| | *CCL3* | 2.5 | 3.8 | 3.7 | 3.1 | | | # 0.07 | 5.6 | 4.4 | 0.012 | | # 0.096 |
| | *CXCL9* | 3.9 | 5.5 | 7.0 | 3.8 | | | # 0.07 | 7.1 | 6.2 | | | |
| Th2 | *IL4* | 44.5 | 93.3 | 40.5 | 88.7 | | 0.003 | | 51.2 | 105.9 | | 0.013 | |
| | *IL13* | 5.5 | 35.4 | 9.6 | 19.0 | | 0.005 | | 5.5 | 38.9 | | 0.001 | |
| | *ARG1* | 11.4 | 61.9 | 9.7 | 76.4 | | 0.007 | | 15.2 | 23.0 | | | 0.026 |
| | *CCL17* | 17.5 | 93.1 | 14.6 | 86.4 | | 0.003 | | 9.3 | 88.8 | | 0.001 | |
| | *CCL26* | 2.2 | 9.3 | 3.1 | 3.9 | | | 0.033 | 2.7 | 6.6 | | 0.001 | |
| Treg | *TGFB1* | 3.1 | 5.7 | 4.8 | 5.5 | | # 0.079 | | 4.5 | 6.5 | | 0.011 | |
| Epithelial cell barrier and mucosal immune function | *IL4R* | 5.7 | 17.1 | 9.4 | 6.7 | | | 0.001 | 8.7 | 10.8 | | 0.032 | |
| | *DCLK1* | 12.1 | 49.3 | 23.2 | 16.7 | | | 0.009 | 25.1 | 31.1 | | 0.019 | |
| | *TSLP* | 235.8 | 600.0 | 367.2 | 472.5 | | # 0.074 | | 433.2 | 483.6 | | 0.042 | |
| | *FFAR2* | 38.9 | 215.7 | 58.2 | 79.1 | | | 0.037 | 52.2 | 140.7 | | 0.017 | |
| | *HDAC2* | 3.5 | 3.6 | 3.1 | 2.5 | # 0.062 | | | 3.0 | 2.8 | # 0.066 | | |
| | *HDAC6* | 6.5 | 10.2 | 9.3 | 5.4 | | | 0.013 | 7.9 | 7.3 | | | |
| | *HDAC9* | 2.6 | 6.1 | 4.3 | 3.7 | | | 0.019 | 4.5 | 6.5 | | 0.012 | |

| | | | | | | | | | | | | | |
|---|---|---|---|---|---|---|---|---|---|---|---|---|---|
| | *RETNLB* | 9.4 | 51.9 | 6.8 | 11.3 | 0.031 | 0.014 | | 7.5 | 24.8 | | 0.004 | |
| Innate immune defence | *IL6* | 4.6 | 15.9 | 5.1 | 8.9 | | 0.02 | | 6.2 | 8.9 | | 0.04 | |
| | *C3* | 2.6 | 3.7 | 2.5 | 3.1 | | 0.01 | | 2.6 | 2.9 | | # 0.064 | |
| | *CD14* | 33.7 | 73.1 | 57.2 | 47.3 | | | 0.094 | 43.3 | 66.1 | | | |
| | *CTLA4* | 2.0 | 3.9 | 3.4 | 2.6 | | | 0.042 | 3.6 | 2.8 | | | # 0.054 |
| | *CXCL10* | 3.8 | 4.0 | 6.7 | 5.3 | 0.04 | | | 6.4 | 4.7 | # 0.059 | | |
| | *PLA2G4A* | 4.2 | 5.1 | 5.7 | 3.7 | | | 0.036 | 4.8 | 4.5 | | | |
| | *PTGES* | 14.9 | 22.3 | 28.6 | 15.8 | | | 0.008 | 16.0 | 18.9 | | | |
| | *PTGS2* | 3.5 | 15.1 | 10.0 | 13.3 | | 0.017 | | 7.0 | 11.6 | | 0.022 | |

**Supplementary Table 5.** Primers used for qPCR.

| Gene | Forward Primer (5' – 3') | Reverse Primer (5' – 3') | Amplicon Length |
|---|---|---|---|
| *IL1A* | TGTGCTAAATAACCTGGATGAGG | GGTTCGTCTTCGTTTTGAGC | 135 |
| *IL1B* | CCAAAGAGGGACATGGAGAA | GGGCTTTTGTTCTGCTTGAG | 123 |
| *IL8* | GAAGAGAACTGAGAAGCAACAACA | TTGTGTTGGCATCTTTACTGAGA | 99 |

| | | | |
|---|---|---|---|
| *IL12B* | GACCAGAAAGAGCCCAAAAAC | AGGTGAAACGTCCGGAGTAA | 70 |
| *IL15* | CGTCATTTTGCAAGAGTCCA | TGGACGATAAACTGCTGTTTGC | 86 |
| *IL17A* | GAGGTACCCCTCCGTGATCT | CTTCCTTCCCTTCAGCATTG | 71 |
| *IFNG* | :CCATTCAAAGGAGCATGGAT | TTCAGTTTCCCAGAGCTACCA | 76 |
| *TNF* | CCCCCAGAAGGAAGAGTTTC | CGGGCTTATCTGAGGTTTGA | 92 |
| *TLR2* | CGGAGGTTGCATATTCCACAG | TGTGAAAGGGAACAGGGAAC | 128 |
| *TLR3* | ATTGTGCAAAAGATTCAAGGTG | TCTTCGCAAACAGAGTGCAT | 130 |
| *TLR4* | TGGTGTCCCAGCACTTCATA | CAACTTCTGCAGGACGATGA | 116 |
| *TLR7* | AGAAGCCCCTTCAGAAGTCC | GGTGAGCCTGTGGATTTGTT | 93 |
| *CD40* | TGAGAGCCCTGGTGGTTATC | GCTCCTTGGTCACCTTTCTG | 90 |
| *CD86* | CATCGTCTGTGTCCTGCAAC | CACAGGTGGCTTTGCATCTA | 82 |
| *CD163* | CACATGTGCCAACAAAATAAGAC | CACCACCTGAGCATCTTCAA | 130 |
| *PRF1* | CTATGGCTGGGACGATGACC | CATGGTTCAAGGCGCACATC | 86 |
| *GZMA* | AAGGGGATCTTCAGCTGCTT | GGGGTTCGACATCTTTTCCT | 99 |
| *GZMB* | CCAGGACCAGGATAATCGAA | GGGTGACGTTGATTGAGCTT | 101 |
| *KLRK* | GATGGTTCCATCCTCTCACC | TGAGCCATAGACTGCACAGC | 75 |
| *INOS* | CAGCCCAAGGTCTATGTTCAAG | ATAGAGGTGGCCTTGCTCCT | 90 |
| *CCL3* | CTCTGCAGCCAGGTCTTCTC | CTACGAATTTGCGAGGAAGC | 97 |
| *CXCL9* | AGCAGTGTTGCCTTGCTTTT | ATGCAGGAACAACGTCCATT | 92 |
| *IL4* | GCAAACATGACCTGTTCTGTG | GCTTCAACACTTTGAGTATTTCTCC | 105 |
| *IL5* | GGGGAAAGATGGAGAGTAACG | CTTTCCATTGTCCACTCGGTA | 83 |
| *IL13* | CCAAGCGAGCAAGTTCCTG | AACTACCCGTGGCGAAAAAT | 110 |
| *ARG1* | TCCAAGGTCTGTGGGAAAAG | ATCGCCATACTGTGGTCTCC | 108 |
| *CCL17* | GGGTGGTACCAGACCTCAGA | GTCCTTGGGGTCAGAACAGA | 90 |
| *CCL22* | CCCTGCGTGTGGTGAAGTAT | ATCTCTCGGTCCCTCAAGGT | 88 |
| *CCL26* | CTGCTTCCAATACAGCCACA | AGCAGCTGTTCCTGGTGAAT | 74 |
| *CCR4* | GGACCCCTTACAATGTGGTG | GAATGGCGTAGTCCAGGTGT | 96 |
| *IL10* | TACAACAGGGGCTTGCTCTT | GCCAGGAAGATCAGGCAATA | 110 |
| *TGFB1* | TCACCGGGGCTGTATTTAAG | AAGGAAGACCCCAGTCAGGT | 110 |
| *FOXP3* | GAAGGACAGCACCCTTTCAA | AGGAAGTCCTCTGGCTCCTC | 111 |
| *IL25* | TGTGTCCACACTGTGTCAGC | GAAGACGGTCTGGTTGTGGT | 89 |
| *IL4R* | CAGAGCTGCCTGCTGTCAT | CTCTCCGGGATCTGAGGACT | 80 |
| *IL13RA1* | TCCCTCCAATTCCTGATCCT | TCCAGTGCAGGGTATCATCA | 75 |
| *DCLK1* | TAAGGCGCAGAGATACAGCA | GGTTCGGTAGAAGCTGCAAT | 85 |
| *TSLP* | ACTAAGGCTGCATTCGCACT | TTTTCCTCATTGCCTGGGTA | 76 |
| *FFAR2* | GCTTCGGGCCCTATAACATA | GCGTTGAGGGAGCTGAATAC | 97 |
| *HDAC1* | GGATCGGTTAGGTTGCTTCA | CCTCCCAGCATCAACATAGG | 96 |
| *HDAC2* | TGCAGTTCATGAAGACAGTGG | CACGCTATCCGTTTGTCTGA | 87 |
| *HDAC3* | GCTGCTGGACGTATGAGACA | GTCTGGATGGAGCGTGAAGT | 110 |
| *HDAC6* | CCCAAATCCATCGCAGATAC | GGCGAACGACTTAGAACTGG | 86 |
| *HDAC9* | GAACAGATGCGACAGCAAAA | CTTTTGTTGCCAAGGGAGAC | 76 |
| *MCT1* | CCGACTTCTGGCAAAAGAAC | GGCTTCTCAGCAGCGTCTAT | 90 |
| *MUC1* | GGATTTCTGAATTGTTTTTGCAG | ACTGTCTTGGAAGGCCAGAA | 116 |
| *MUC2* | GCACGTCTGCAACAAGGAC | CAAAGCCCTCCAGGCAGT | 125 |
| *RETNLB* | TCCCTCTGCTCCAAGAAAGA | CAAGCACAGCCAGTGACAAC | 99 |
| *SLC2A5* | GGTCATCTCCACCATCATCC | GCGCTCAGGTAGATCTGGTC | 90 |
| *SLC5A1* | TCTCATGAGCTCCCTGACCT | CTCTCTTCCGGATCTTGGTG | 83 |
| *SLC5A8* | TGGGACAAATTGGATGACAA | CCATCAGTGGAGTCCTTTCAA | 86 |
| *TFF2* | GCTGCTTCGACTCCCAAGT | CATGACGCACTCCTCAGACT | 80 |
| *TFF3* | TGTTCTGGCTGCTAGTGGTG | CAGTCCACCCTGTCCTTGG | 112 |
| *IL6* | TGGGTTCAATCAGGAGACCT | CAGCCTCGACATTTCCCTTA | 116 |

| | | | |
|---|---|---|---|
| *IL18* | CAATTGCATCAGCTTTGTGG | TCCAGGTCCTCATCGTTTTC | 78 |
| *CXCL10* | CCCACATGTTGAGATCATTGC | GCTTCTCTCTGTGTTCGAGGA | 141 |
| *C3* | ATCAAATCAGGCTCCGATGA | GGGCTTCTCTGCATTTGATG | 76 |
| *CD14* | GGGTTCCTGCTCAGATTCTG | CCCACGACACATTACGGAGT | 164 |
| *CLDN3* | ATCGGCAGCAGCATTATCAC | ACACTTTGCACTGCATCTGG | 94 |
| *CTLA4* | CTCCTGTACCCACCACCCTA | AGAATCTGGGCATGGTTCTG | 84 |
| *DEFB1* | TTCCTCCTCATGGTCCTGTT | CATCTTTGGAGCACACTTGC | 114 |
| *OCLN* | GACGAGCTGGAGGAAGACTG | GTACTCCTGCAGGCCACTGT | 102 |
| *PLA2G4A* | CGTACCCCTTGATCCTGAGA | CTTGGCCTTGCAGAAAAGTC | 73 |
| *PTGES* | TGTACGTAGTGGCCATCATCA | CTCCGTGTCTCTGAGCATCC | 84 |
| *PTGS2* | GAACTTACAGGAGAGAAGGAAATGG | TTTCTACCAGAAGGGCAGGA | 94 |
| *SAA* | GCTAAAGTGATCAGCGATGC | AGTGGTTGGGGTCCTTGC | 145 |
| **Housekeeping genes** | | | |
| *GAPDH* | ACCCAGAAGACTGTGGATGG | AAGCAGGGATGATGTTCTGG | 79 |
| *RLP13A* | ATTGTGGCCAAGCAGGTACT | AATTGCCAGAAATGTTGATGC | 76 |
| *PPIA* | CAAGACTGAGTGGTTGGATGG | TGTCCACAGTCAGCAATGGT | 138 |
